# Resonance in physiologically structured population models

**DOI:** 10.1101/2021.01.18.427157

**Authors:** Kevin Gross, André M. de Roos

## Abstract

Ecologists have long sought to understand how the dynamics of natural populations are affected by the environmental variation those populations experience. A transfer function is a useful tool for this purpose, as it uses linearization theory to show how the frequency spectrum of the fluctuations in a population’s abundance relates to the frequency spectrum of environmental variation. Here, we show how to derive and to compute the transfer function for a continuous-time model of a population that is structured by a continuous individual-level state variable such as size. To illustrate, we derive, compute, and analyze the transfer function for a size-structured population model of stony corals with open recruitment, parameterized for a common Indo-Pacific coral species complex. This analysis identifies a sharp multi-decade resonance driven by space competition between existing coral colonies and incoming recruits. The resonant frequency is most strongly determined by the rate at which colonies grow, and the potential for resonant oscillations is greatest when colony growth is only weakly density-dependent. While these resonant oscillations are unlikely to be a predominant dynamical feature of degraded reefs, they suggest dynamical possibilities for marine invertebrates in more pristine waters. The size-structured model that we analyze is a leading example of a broader class of physiologically structured population models, and the methods we present should apply to a wide variety of models in this class.

## 1 Introduction

Ecologists have long sought to understand how environmental variation affects the dynamics of populations in nature (e.g., Ives, 1995; Higgins et al., 1997; Bjørnstad and Grenfell, 2001; Greenman and Benton, 2003, among many others). One way to build this understanding is to compare the frequency spectrum (also called the power spectrum, or just spectrum) of the fluctuations in a population’s abundance to the frequency spectrum of the environmental fluctuations that act upon the population (Nisbet and Gurney, 1982; Greenman and Benton, 2005a). Comparing these spectra reveals how the endogenous processes acting within a population amplify, muffle, or otherwise translate environmental fluctuations into oscillations in the population’s dynamics.

One useful tool for comparing spectra in population models is a *transfer function*, whose use in population ecology was pioneered by Nisbet & Gurney (Gurney and Nisbet, 1980; Nisbet and Gurney, 1982). To calculate a transfer function, one first constructs a model for the population dynamics that depends on one or more time-varying environmental drivers and converges to a locally stable equilibrium when the environment is constant. The model is linearized about its constant-environment equilibrium, and the linearized model is transformed to the frequency domain. Solving the resulting system of equations yields an expression for the Fourier transform of the population fluctuations as a linear function of the Fourier transform(s) of the environmental driver(s). The frequency-dependent proportionality between the two spectra gives the transfer function. Transfer functions are especially useful for characterizing resonances in population dynamics, in which endogeneous feedback within the population predisposes the population to oscillate at a particular harmonic frequency in suitably perturbed environments. Transfer functions and related linearization approaches have since been used to study, for example, models of infectious disease (Ruxton, 1996), fisheries (Bjørnstad et al., 2004; Worden et al., 2010), and simple food webs (Ripa et al., 1998).

In this paper, we provide new methods to derive and to compute transfer functions for a size-structured population model in continuous time. This model is a leading example of a broader class of physiologically structured, continuous-time population models (PSPMs) in which individuals are classified by one or more continuous state variables such as size. PSPMs provide a natural modeling framework for analyzing non-linear dynamical phenomena that result from size- or stage-mediated interactions, such as resource competition between juveniles and adults (ten Brink et al., 2019), size-dependent maturation (de Roos and Persson, 2013), and cannibalism (Claessen et al., 2000). The now-mature theory of PSPMs is described in Metz and Diekmann (1986) and Diekmann et al. (2007), while de Roos (1997) provides a gentler introduction. We conjecture that the methods we describe here can be extended to many other PSPMs.

Our work is motivated by a size-structured model of the common Indo-Pacific reef-forming coral species complex *Pocillopora verrucosa*. Previously, we used this model to investigate coral population dynamics in strongly disturbed waters (Hall et al., 2021). Our analysis there found that, in a disturbance-free environment, coral cover exhibited under-damped dynamics that eventually converged to a stable equilibrium. The underdamped dynamics suggest that environmental stochasticity could drive regular oscillations in total coral cover in moderately disturbed environments (Nisbet and Gurney, 1976), and simulations confirm that this is so (Fig. 1). Here, we compute transfer functions to characterize the resonant frequency in these dynamics, and study how resonance depends on the population’s underlying vital rates.

**Figure 1:**
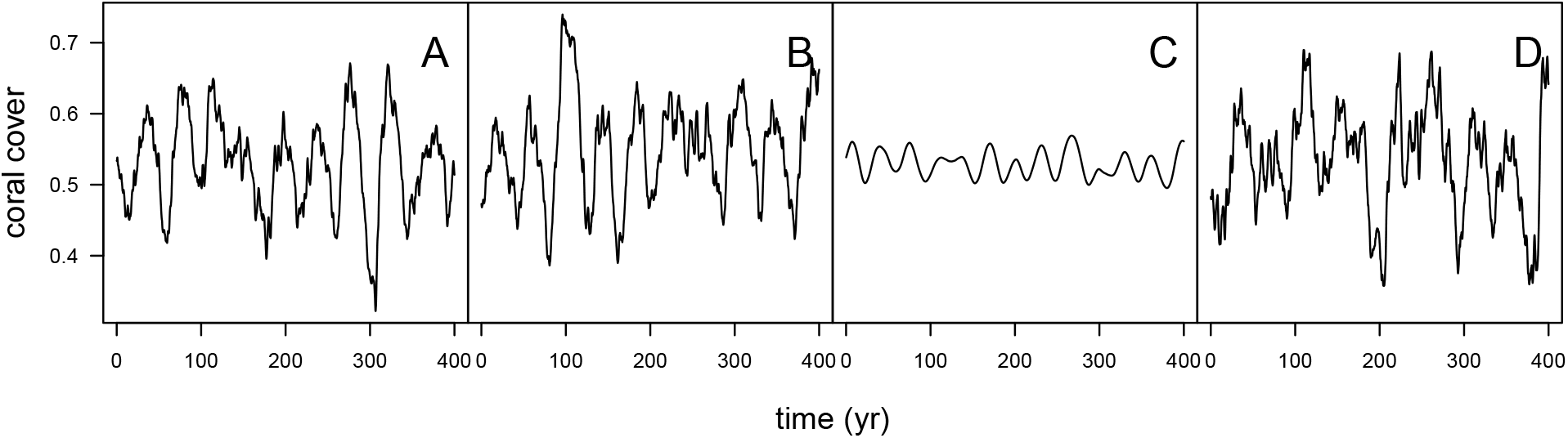
Environmental stochasticity excites quasi-cycles in the *Pocillopora verrucosa* model. Panels (A)–(C) show 400 yr of simulated dynamics when (A) colony growth, (B) colony mortality, or (C) recruitment fluctuates in response to environmental variation, while panel (D) shows dynamics when all three vital rates fluctuate concurrently and independently. When the environment is held constant at its time-averaged value, coral cover converges to a steady state of ≈ 0.525. (Coral cover is measured as a proportion of available substrate, and is thus dimensionless.) Simulations were initiated at this steady state, and the first 200 yr were discarded to eliminate transients. More details about the simulation methods are given in §3.2.

More broadly, this model contributes to a rich literature on size-structured population models for openly recruiting populations of benthic marine invertebrates. A prevailing theme to emerge from this literature is that intraspecific space competition creates delayed density dependence that promotes population oscillations, especially when available substrate is scarce (Roughgarden et al., 1985). Essentially, established individuals and incoming recruits all compete for, and are limited by, available substrate. Thus, the growth of existing colonies interferes with the recruitment of new colonies that will comprise the bulk of the population some time hence. In deterministic models, the resulting population oscillations can manifest either as stable population cycles or as decaying oscillations en route to a stable point equilibrium. The mathematical study of these populations was pioneered by Roughgarden et al. (1985), and has been continued by Bence and Nisbet (1989), Pascual and Caswell (1991), Muko et al. (2001), and Artzy-Randrup et al. (2007).

The rest of this paper proceeds as follows. First, we present a size-structured population model with environmental forcing. The unforced version of this model derives from the classic McKendrick - von Foerster model (McKendrick, 1925; Von Foerster, 1959), and a more detailed presentation of its construction can be found in, e.g., de Roos (1997). We then derive the transfer functions for this model. To illustrate, we then calculate the transfer functions for our *P. verrucosa* model. Subsequent numerical explorations show how the resonant oscillations in the coral model are affected by changes in vital rates, density-dependent growth, and self-seeding. The appendix describes how to compute the transfer functions, while the supplementary material provides additional figures and results as well as R computer code (R Core Team, 2018) to reproduce all primary results.

## 2 Mathematical development

### 2.1 A size-structured population model with environmentally forced vital rates

We develop our results in the context of a continuous-time model for a size-structured population. Use 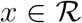 to denote the size of an individual in this population. We assume that size is the lone individual-level state variable, meaning that individuals of the same size at the same time will experience the same demographic fate. Let *x*_0_ denote the size of new individual entering the population, and let *x*_1_ > *x*_0_ denote the maximum size an individual can attain. The population state is described by the size distribution of individuals at time *t*, denoted *n*(*t*, *x*), such that *n*(*t*, *x*) *dx* is the population density of individuals with sizes in (*x*, *x* + *dx*) at time *t*.

We assume that an individual’s demography depends on *n*(*t, x*) only through the total population size *C*(*t*). (We use *C*(*t*) in anticipation of the coral model to follow.) Let *A*(*x*) be the contribution of a size *x* individual to the total population size. In the coral model, *A*(*x*) will be the planar area of a coral colony. Then we have simply

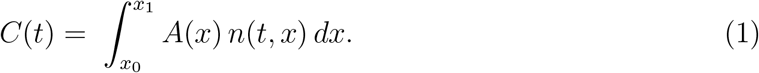

A size-structured population model requires specifying three demographic vital rates: growth, mortality, and fecundity. We assume that each of these rates may depend on an individual’s size *x*, the total population size *C*(*t*), and a separate, time-varying environmental input for each rate. Use *g*(*x*, *C*(*t*), *E*_1_(*t*)) to denote the growth rate of a size *x* individual when the total population size is *C*(*t*) and the environmental input affecting growth is *E*_1_(*t*). The growth rate is simply the time derivative of an individual’s size with respect to time. Similarly, use *μ*(*x*, *C*(*t*), *E*_2_(*t*)) and *β*(*x*, *C*(*t*), *E*_3_(*t*)) to denote the mortality and fecundity rates of a size *x* individual, respectively, when the total population size is *C*(*t*). The environmental inputs affecting mortality and fecundity are *E*_2_(*t*) and *E*_3_(*t*), respectively. The fecundity rate is the rate at which an individual parent contributes new offspring to the population.

Together with eq. 1, assembling the components yields the full model (see de Roos (1997) for details)

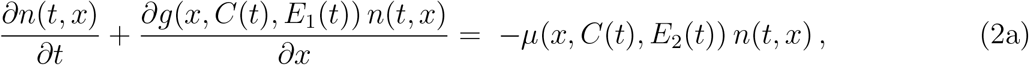

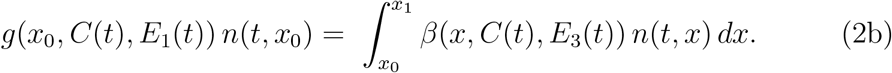

Eq. 2 is a size-structured, non-linear, environmentally forced version of the McKendrick - von Foerster model (McKendrick, 1925; Von Foerster, 1959), where eq. 2a is a conservation equation and eq. 2b is a boundary condition.

### 2.2 Derivation of the transfer function for a size-structured PSPM

The main contribution of this paper is a method to derive and compute the transfer functions for the model in eq. 1 – 2. Before embarking on the derivation, we preview it briefly to make its destination clear. We require that the environmental fluctuations are generated by a stationary process, and we use 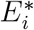 to denote the time-averaged values of the environmental variables *E_i_*(*t*) for *i* = 1, 2, 3. We also require that the total population size *C*(*t*) approaches a locally stable equilibrium *C*\* when the *E_i_*(*t*) are held constant at 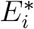. The model is linearized about its constant-environment equilibrium, with deviations of *C*(*t*) and *E_i_*(*t*) from equilibrium defined as

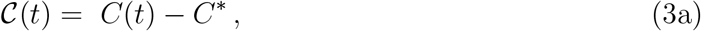

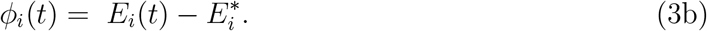

Now define the Fourier transforms of 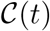 and *ϕ_i_*(*t*) as 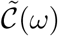 and 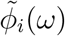, respectively, where *ω* is the angular frequency and we recall that a Fourier transform is defined as, for example,

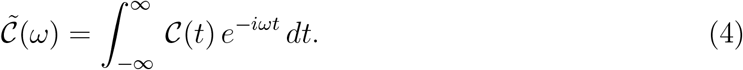

Ultimately, we seek an expression for 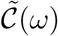 in the form

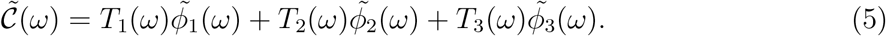

The term *T_i_*(*ω*) is the transfer function for the environmental driver *E_i_*(*t*), as it “transfers” the environmental spectrum 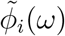 to the population spectrum 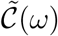. The derivation of the transfer functions proceeds in three steps. First, the “Murphy trick” (Murphy, 1983; Metz and Diekmann, 1986) is used to reformulate the model in terms of an individual’s age. Second, the reformulated model is linearized about its equilibrium and then transformed to the spectral domain using standard techniques. Third, the resulting system of equations is solved to yield an expression in the form of eq. 5. This concludes the preview.

We first use the “Murphy trick” (Murphy, 1983; Metz and Diekmann, 1986) to re-formulate the model in eq. 1 – 2 in terms of an age-distribution *m*(*t*, *a*) and a size-at-age function *x*(*t*, *a*) for individuals in the population. Here, *m*(*t*, *a*) *da* gives the abundance of individuals at time *t* with ages in the interval (*a*, *a* + *da*) and *x*(*t*, *a*) gives the size of age *a* individuals at time *t*. The reformulated model is

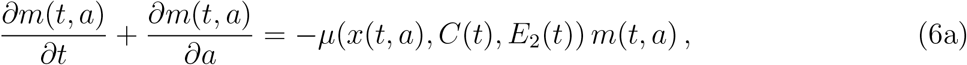

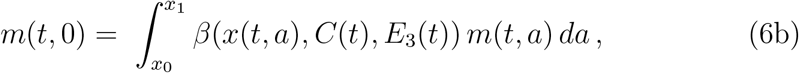

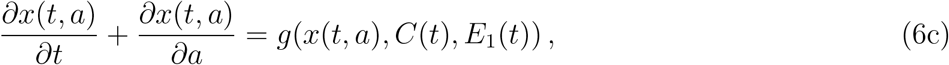

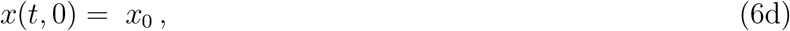

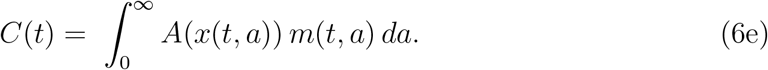

Next, define the “birth rate” 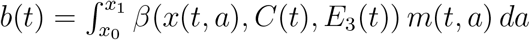 as the rate at which new individuals enter the population. Also define 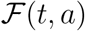 as the survival of individuals recruited at time *t* − *a* to age *a*; we call 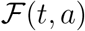 the survival function. Writing 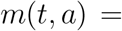 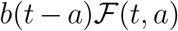 allows the model to be rewritten in terms of PDEs for *x*(*t*, *a*) and 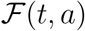 and a renewal equation for *b*(*t*), leading to

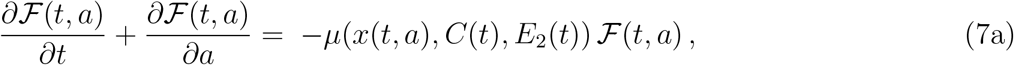

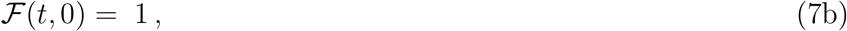

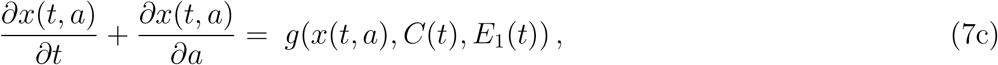

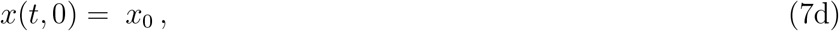

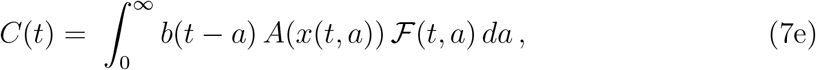

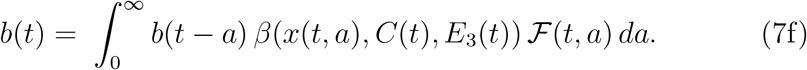

We next linearize eq. 7 about its equilibrium in the usual way. Let 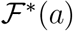, *x*\*(*a*), *C*\*, *b*\*, and 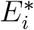 denote the equilibrium values of 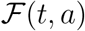, *x*(*t*, *a*), *C*(*t*), *b*(*t*), and *E_i_*(*t*), respectively. Write the equilibrium growth-at-age, mortality-at-age, fecundity-at-age, and area-at-age relationships as *g*\*(*a*) = *g*(*x*\*(*a*), *C*\*, 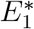, *μ*\*(*a*) = *μ*(*x*\*(*a*), *C*\*, 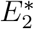), *β*\*(*a*) = *β*(*x*\*(*a*), *C*\*, 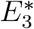), and *A*\*(*a*) = *A*(*x*\*(*a*)), respectively. Define the following deviations from equilibrium, along with those in eq. 3:

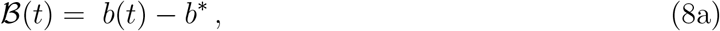

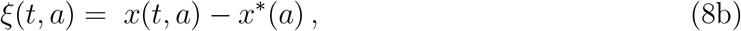

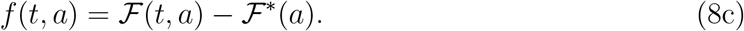

Use 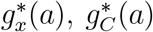, and 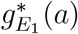 to denote the partial derivative of *g*(*x*, *C*, *E*_1_) with respect to *x*, *C*, and *E*_1_, respectively, evaluated at age *a* and at equilibrium. That is, for example,

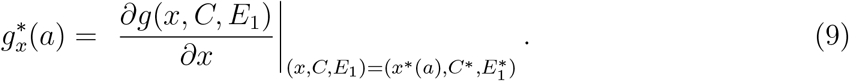

Denote the equivalent partial derivatives of mortality (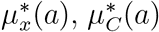, and 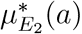) and fecundity (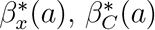, and 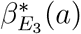) similarly. Also use 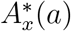 to denote the derivative *dA*(*x*) / *dx* evaluated at *x* = *x*\*(*a*). Linearization thus yields the following first-order approximation of eq. 7 in the neighborhood of the equilibrium

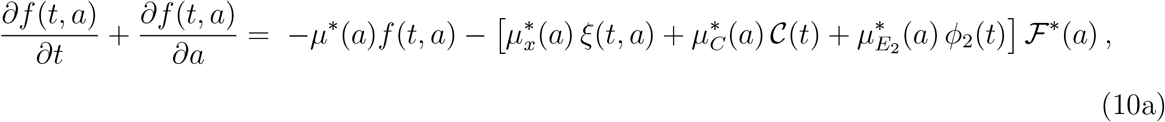

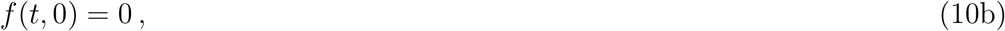

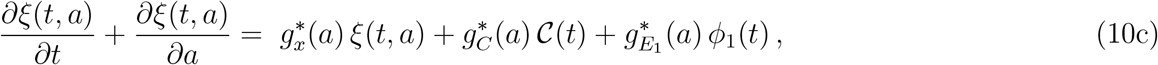

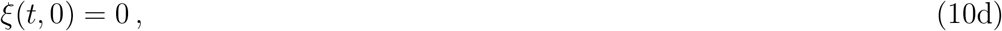

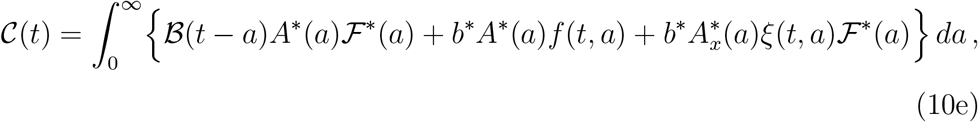

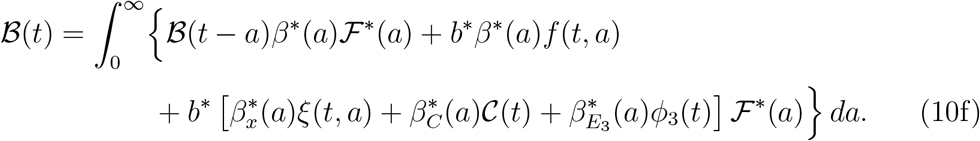

Next, let 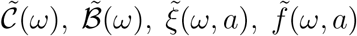, and 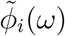 be the Fourier transforms of 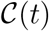, 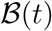, *ξ*(*t*, *a*), *f* (*t*, *a*), and *ϕ_i_*(*t*), respectively. Taking the Fourier transformation of eq. 10 then yields

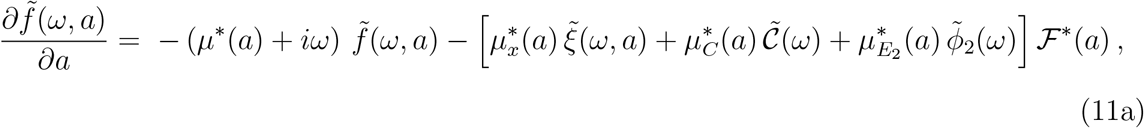

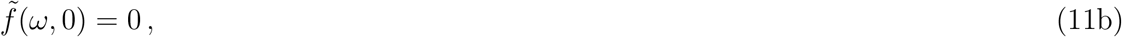

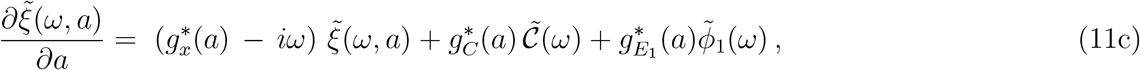

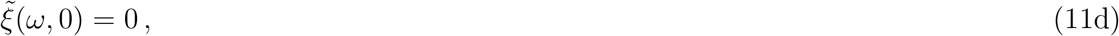

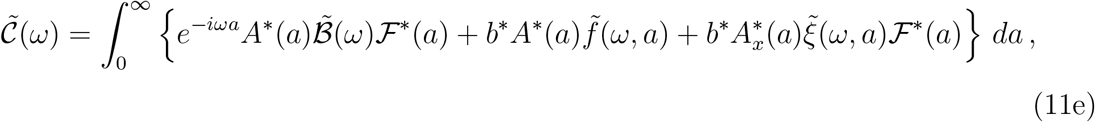

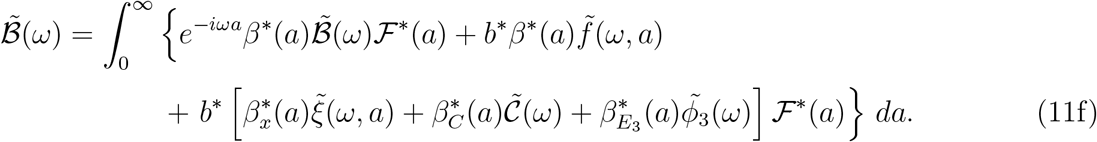

Finally, we solve for 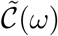 in terms of 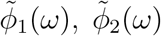, and 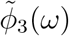 by following a path similar to the appendix of Gurney and Nisbet (1980). To begin, note that in eq. 11c holding *ω* constant gives is a linear differential equation for 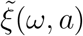. Thus we can use an integrating factor to solve for 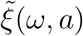 as

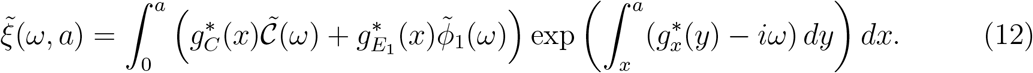

Now factor 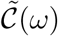 and 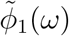 out of the integral to obtain

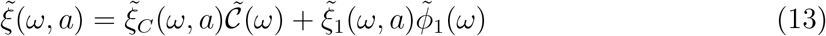

with

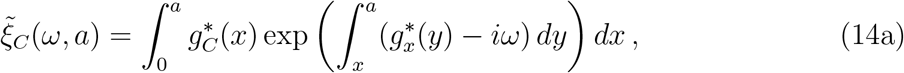

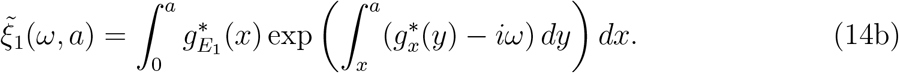

Next, plugging the solution for 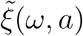 into eq. 11a allows us to re-write eq. 11a as

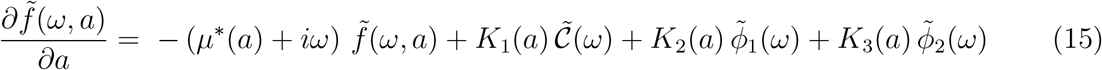

with

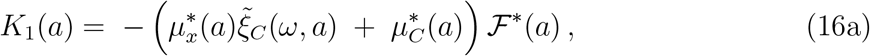

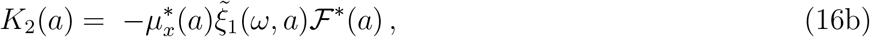

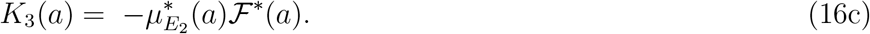

This can also be solved with an integrating factor to obtain

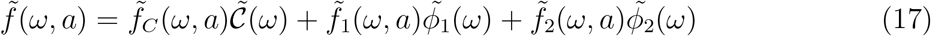

in which

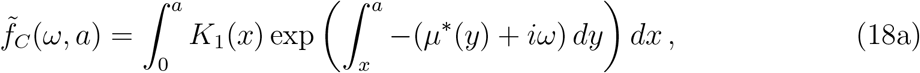

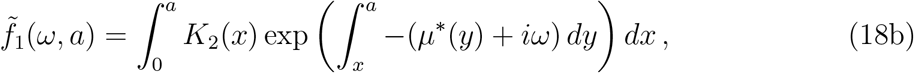

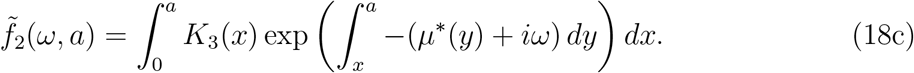

Finally, plugging solutions for 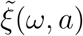 and 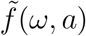 into eq. 11e – 11f gives, after some algebra,

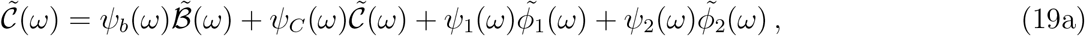

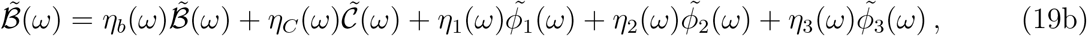

where

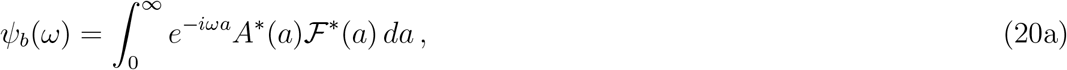

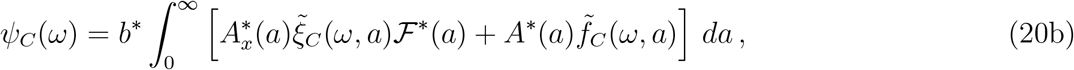

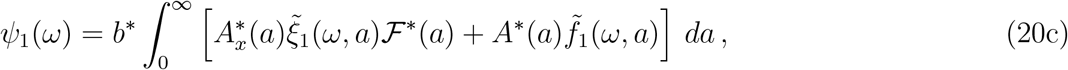

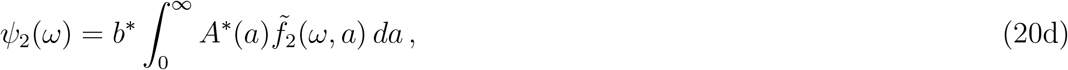

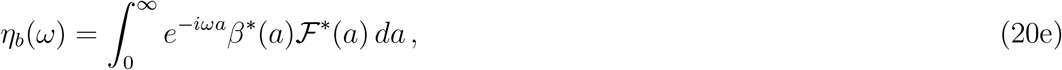

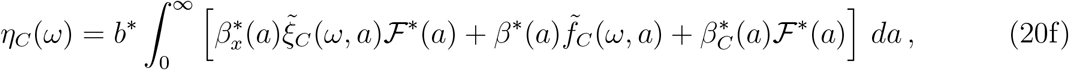

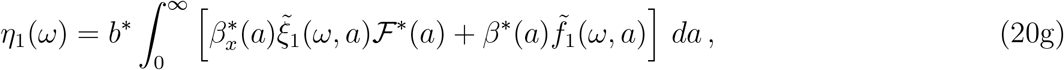

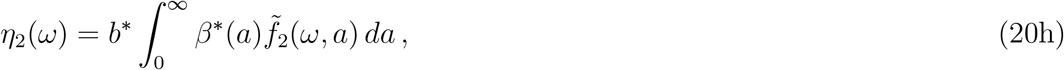

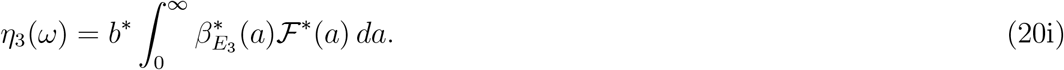

Note that eq. 19 provides a linear system of equations that show implicitly how fluctuations in the environmental drivers combine to generate fluctuations in the model’s two fundamental variables, total population size *C*(*t*) and total population birth rate *b*(*t*). The integrals in eq. 20 are merely the coefficients in this linear system. Solving eq. 19 for 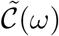 thus yields an expression in the form of eq. 5, with

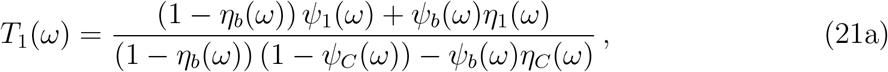

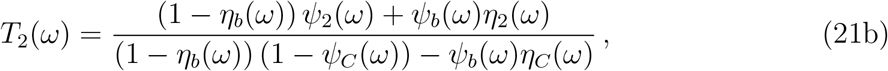

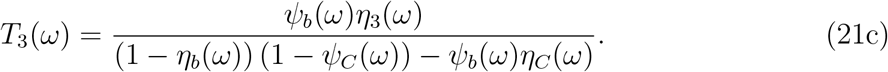

Eq. 21 provides the transfer functions that we seek.

While the formulae for the transfer functions look opaque, they have a noteworthy structure. Each transfer function is a ratio, and those ratios share a common denominator. Moreover, the terms that capture how the vital rates depend on environmental variation, 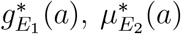, and 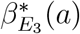, only affect the numerators of the transfer functions, while the terms capturing the density-dependence, 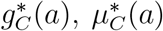, and 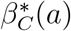, only affect the denominator. Terms that describe the size-dependence of vital rates affect both the numerator and denominator. This structure parallels the structure of Bjørnstad et al. (2004)’s transfer functions for discrete-time, age-structured models. There, as Bjørnstad et al. (2004) observed, the transfer functions also took the form of ratios, with numerators that integrated the effects of environmental stochasticity, and denominators that integrated the effects of direct and delayed density dependence.

Finally, because both 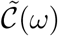 and 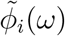 scale with *C*\* and 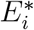, we find it more useful to compute a scaled transfer functions *T_i,s_*(*ω*) that satisfy

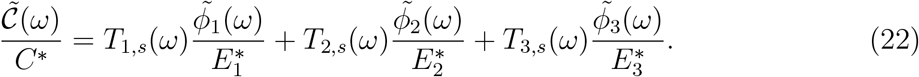

Simple algebra shows that

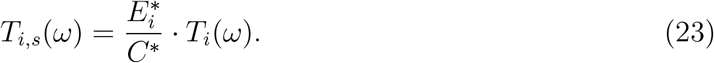

The appendix describes how to calculate the transfer functions by approximating the integrals in eq. 20 with the numerical solution of a system of coupled ODEs (Kirkilionis et al., 2001).

## 3 Application

### 3.1 Coral model

We illustrate this method by calculating the transfer functions for a size-structured population model of the common Indo-Pacific coral species complex *Pocillopora verrucosa* as it occurs at 10m depth on the north shore of Mo’orea, French Polynesia. More details about the model and its parameterization can be found in Hall et al. (2021).

Stony corals are colonial organisms. We count a colony as a discrete individual because individual polyps in the same colony are tightly coupled physiologically (Mackie, 1986). Size is a natural state variable for coral colonies, because the colony demography is thought to be determined by the colony’s size (Hughes, 1984; Madin et al., 2014), and because colony size is more easily measured than colony age. We quantify the size of a colony by its effective diameter, such that the planar area of a diameter *x* colony is *A*(*x*) = *π*(*x*/2)^2^.

Let *n*(*t*, *x*) give the size distribution of the population at time *t* in units of colonies per *m*^2^. Coral population abundance is usually measured by the proportion of available habitat covered by live colonies. Following this practice, *C*(*t*) in the model (eq. 1) gives the proportion of substrate covered by live coral. We will define demographic inputs so that both colony growth and recruitment are 0 when *C*(*t*) = 1, thus ensuring that *C*(*t*) ≤ 1. The proportion of available substrate is then just 1 − *C*(*t*).

For *P. verrucosa*, newly settled spat have a diameter of *x*_0_ = 0.4 mm (Babcock, 1991), and mature coral colonies have a maximum diameter of *x*_1_ = 0.5 m (Veron, 2000). We model colony growth by setting *g*(*x, C*(*t*), *E*_1_(*t*)) = *g*_0_(*x*) × *g*_1_(*C*(*t*)) × *E*_1_(*t*), where *g*_0_(*x*) captures the size-dependence of growth, *g*_1_(*C*(*t*)) captures density-dependence, and the environmental driver has an average value of 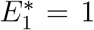. We estimate *g*_0_(*x*) from annual photoquadrat monitoring of apparent *P. verrucosa* colonies from 2011–18 (P. J. Edmunds, personal communication). Our fit suggests that *g*_0_(*x*) takes a quadratic, concave form, constrained so that colonies stop growing when they reach their maximum size (Fig. 2A). Because the reef from which these data were obtained was recovering from a large die-off in 2002–10 (Kayal et al., 2012; Holbrook et al., 2018), the colonies in the data set are all small (*x* ≤ 0.12 m), and thus our fit requires substantial extrapolation. To capture density dependence, we follow previous modeling work (Muko et al., 2001) and assume that growth is proportional to free space, *g*_1_(*C*(*t*)) = 1 − *C*(*t*). Different forms of density-dependence are explored in §3.3 below.

**Figure 2:**
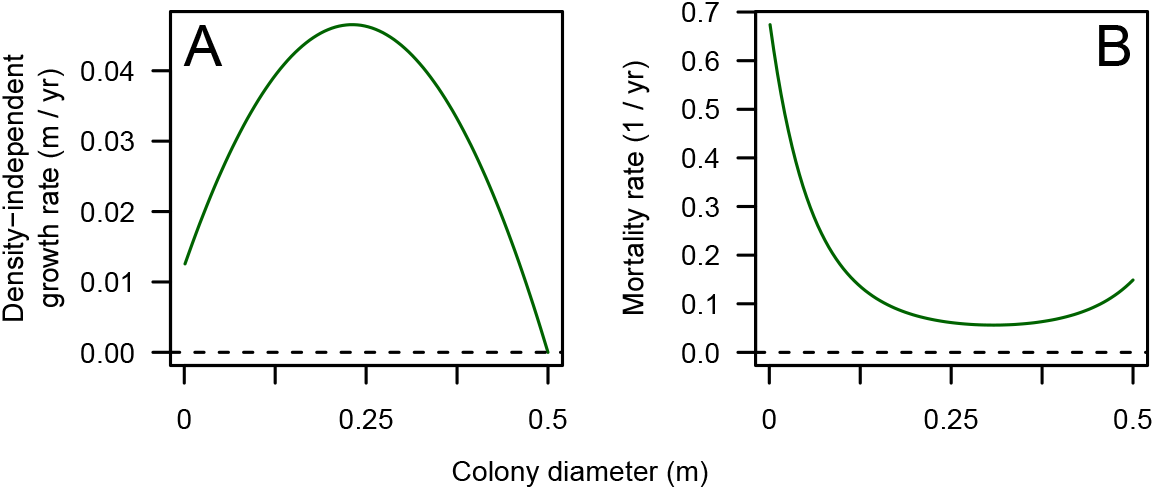
Size-dependent vital rates for the *Pocillopora verrucosa* model. (A): Size-dependent component of growth, *g*_0_(*x*). (B): Size-dependent mortality rate, *μ*(*x*). Redrawn from Hall et al. (2021).

For mortality, we consider only background mortality that acts more or less steadily through time. We do not consider episodic mortality events such as mass bleaching or outbreaks of corallivores that occur as distinct mortality pulses. Following Madin et al. (2014), we estimate a convex relationship between mortality and size, with mortality lowest for intermediate-sized colonies (Fig. 2B). This convex relationship arises because small colonies are most susceptible to overgrowth from space competitors (Ferrari et al., 2012), and large colonies are most susceptible to fragmentation or dislodgement from hydrodynamic stress (Madin et al., 2014). Our mortality curve is parameterized to match the mortality of small colonies observed by Holbrook et al. (2018). Little seems to be known about density-dependence in coral mortality, so we follow previous modeling work (Roughgarden et al., 1985; Artzy-Randrup et al., 2007) in assuming that mortality is independent of population density. Like growth, we assume that the environment acts multiplicatively on mortality, and so we set *μ*(*x, C*(*t*), *E*_2_(*t*)) = *μ*(*x*)*E*_2_(*t*), where *μ*(*x*) captures size-dependent mortality and 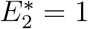.

Like most coral reefs, the coral populations at Mo’orea are thought to recruit primarily by immigration of pelagic larvae spawned elsewhere (Holbrook et al., 2018). Thus, we depart from the closed-population model in §2 and assume instead that immigrating larvae arrive at a rate *s E*_3_(*t*), where *s* gives the average immigration rate and 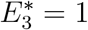. We follow previous modeling work (Roughgarden et al., 1985; Artzy-Randrup et al., 2007) in assuming that the proportion of immigrating larvae that successfully settle equals the proportion of available substrate. Thus, successful recruits enter the population at the rate *s E*_3_(*t*)(1 − *C*(*t*)). This assumption changes the boundary condition of the model (eq. 2b) to

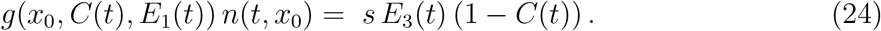

Note that the coral model can be found as a special case of the general model in §2 by setting *β*(*x, C*(*t*), *E*_3_(*t*)) = *s E*_3_(*t*)(1−*C*(*t*))*A*(*x*)/*C*(*t*). This recruitment model also simplifies calculation of the transfer functions, because the “birth rate” *b*(*t*) now depends on the population state only through the population size, *C*(*t*), as *b*(*t*) = *sE*_3_(*t*)(1 − *C*(*t*)). Making this substitution at eq. 7e allows us to replace eqs. 7e–7f with the single equation

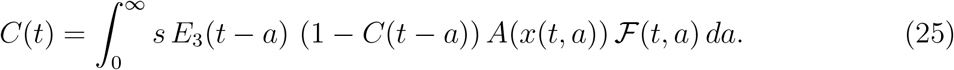

We thus eliminate the need to track *b*(*t*), which simplifies many of the subsequent steps.

We use data from Holbrook et al. (2018) to estimate *s* = 50 colonies m^−2^ yr^−1^. With these parameters, the model gives an average population size in which corals cover slightly more than half of the available substrate, *C*\* ≈ 0.525.

### 3.2 Transfer function

Scaled transfer functions for the *P. verrucosa* model show sharp peaks at a frequency of *ω* / 2*π* ≈ 0.0273 cycles per yr, corresponding to a cycle period of ≈ 36.5 yr (Fig. 3). These sharp peaks indicate that stochastically excited resonance will emerge in *C*(*t*), as long as the spectra of the environmental drivers do not contain sharp peaks themselves. The frequencies at which the peak occurs are not exactly the same for the three transfer functions, but they differ by < 1%. The modulus of the scaled transfer function at the resonant frequency is roughly 3 times greater for growth and mortality than for recruitment, reflecting the greater sensitivity of coral cover to growth and mortality in this model (Hall et al., 2021).

**Figure 3:**
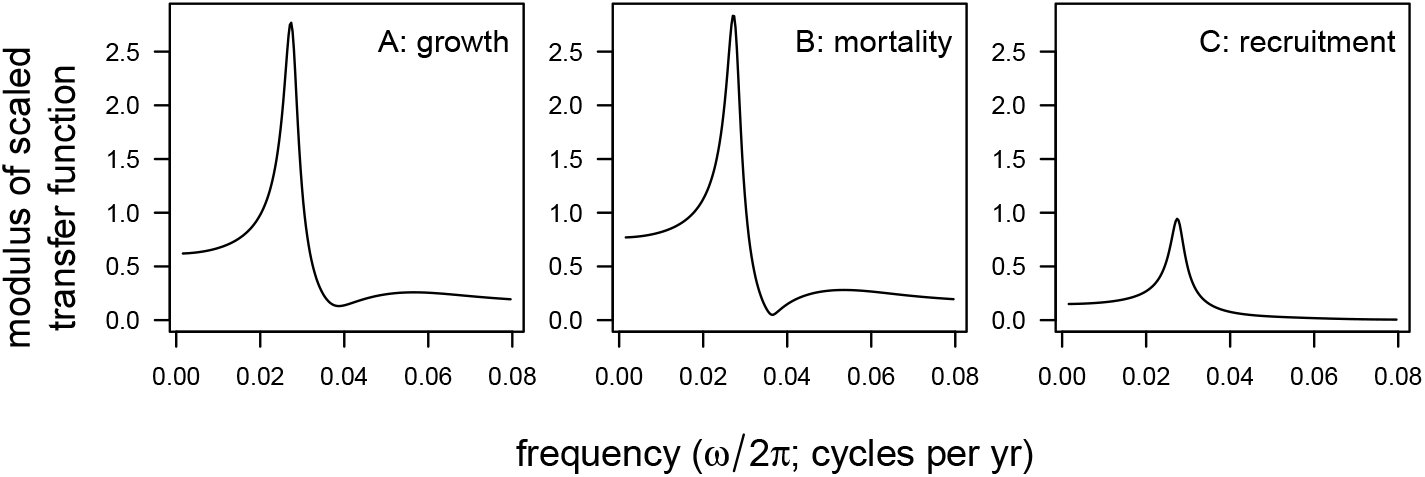
The transfer functions of the *P. verrucosa* model show sharp peaks at a frequency of *ω* / 2*π* ≈ 0.0273 cycles per year, indicating the potential for stochastically excited quasi-cycles. Panels show the modulus of the scaled transfer function for (A) growth, *T*_1,*s*_, (B) mortality, *T*_2,*s*_, and (C) recruitment, *T*_3,*s*_.

To verify this resonance, we used the Escalator Boxcar Train (EBT) method (de Roos, 1988) to simulate dynamics with stochastically forced vital rates (Fig. 1). In brief, the EBT divides the population into a collection of cohorts, where each cohort consists of individuals that enter the population during a time interval of length Δ*t*. A system of coupled ODEs then tracks the abundance of each cohort and the average size of the individuals in each cohort. To add environmental forcing, we allowed either *E*_1_(*t*), *E*_2_(*t*), or *E*_3_(*t*) to be a piecewise constant function with separate values for the time intervals [0, Δ*t*), [Δ*t*, 2Δ*t*), …. Successive values of the forcing variable were determined by a first-order autoregression process with a stationary coefficient of variation of 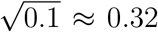, and a correlation between successive values of 0.9. We used a time step of Δ*t* = 0.1 yr. Thus, in these simulations the environmental variation is “pink”, that is, enriched in low-frequency oscillations, as befits environmental fluctuations in nature (Halley, 1996). Simulations clearly show the resonance emerging at the frequency predicted by the transfer functions (Fig. 1), and with much greater sensitivity to fluctuations in growth or mortality than to fluctuations in recruitment. In addition, we fit a smoothed periodogram to longer (5000 yr) simulations, which estimated spectral peaks in the Fourier transform of coral cover at 0.0270, 0.0260, and 0.0282 cycles per yr when growth, mortality, or recruitment fluctuated alone, respectively, and at 0.0254 cyles per yr when all three vital rates fluctuated concurrently.

### 3.3 Numerical explorations

To better understand of how colony vital rates affect resonance in this model, we conducted several numerical experiments. First, we multiplied growth, mortality, and recruitment functions by a factor between 0.5 – 2 and computed the resulting frequency and modulus of the peak in the scaled transfer function. When changing growth and mortality, we scaled recruitment so that equilibrium cover would remain constant. Colony growth has the most straightforward effect on resonance, as faster growth accelerates oscillations by allowing incoming recruits to rebuild cover more quickly during the cycle’s rebound phase (Fig. 4A). However, colony growth has a more complicated effect on the amplitude of resonant cycles, as the strength of resonance peaks at an intermediate growth rate (Fig. 4D). Mortality also has a complicated effect on resonance (Fig 4B,E), perhaps because mortality affects both the high-mortality bottleneck for smaller corals and the clearance of older, larger colonies from the population. Increased recruitment has a small effect, slowing resonant oscillations while increasing their amplitude (Fig 4C,F). Because we did not fix equilibrium cover when varying recruitment, the effect of recruitment on resonance is likely explained by recruitment’s effect on average cover.

**Figure 4:**
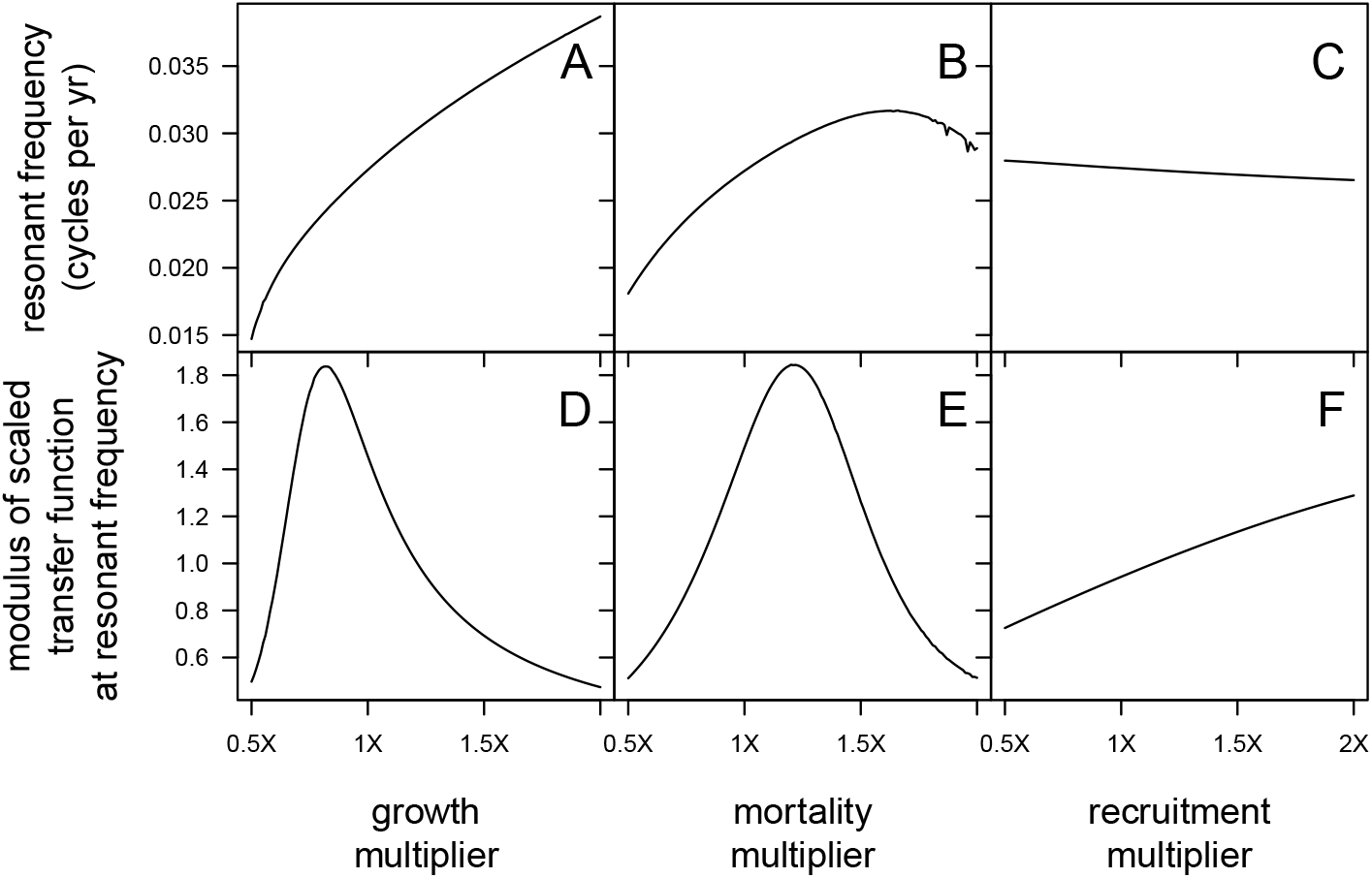
Vital rates have variable effects on resonant frequency and strength in the *P. verrucosa* model. Panels in the top row show how the resonant frequency changes as (A) growth, (B) mortality, and (C) recruitment functions are multiplied by a factor of 0.5 – 2. Panels in the bottom row show how the modulus of the scaled transfer function at the resonant frequency changes as (D) growth, (E) mortality, and (F) recruitment multipliers are varied. In each case, the transfer function considered is the one associated with the varied vital rate, that is, *T*_1,*s*_(*ω*) in (A) and (D), *T*_2,*x*_(*ω*) in (B) and (E), and *T*_3,*s*_(*ω*) in (C) and (F). In (A), (B), (D), and (E), recruitment is tuned to hold equilibrium coral cover constant. In (C) and (F), equilibrium cover varies as recruitment changes.

Second, although there is empirical evidence that colonies grow more slowly in crowded conditions (Tanner, 1997), little is known about exactly how density-dependence affects growth. Our assumption that growth is proportional to free space is motivated by an appeal to parsimony, and precedent from previous modeling work (Muko et al., 2001). In the Supplementary Information, we show that resonant cycles become more frequent and smaller in amplitude as density-dependence in growth becomes stronger (Fig. S1b, c). This agrees with Artzy-Randrup et al. (2007)’s finding that strong density dependence in growth dampens population oscillations by allowing colony growth to counterbalance departures from equilibrium cover more rapidly.

Finally, we conducted a third numerical experiment to investigate how internal recruitment, or “self-seeding”, affects resonance. Empirical evidence for the importance of internal recruitment of corals varies widely (Sammarco and Andrews, 1989; Gilmour et al., 2009; Baskett et al., 2010), while environmental change threatens to disrupt larval connectivity among reefs (Jones et al., 2009). In the Supplementary Information, we show that the frequency of resonance is almost entirely independent whether a population recruits internally or externally (Fig. S3A). However, self-seeding dampens resonant oscillations (Fig. S3B). Closed populations are more strongly buffered against resonant oscillations because increased space competition caused by an excess of coral cover is partially counterbalanced by an overproduction of recruits, whose growth in subsequent years dampens the ensuing oscillation. This result contrasts with models of space competition for unstructured populations (Levins, 1969) in which open recruitment stabilizes population dynamics more strongly than closed recruitment.

## 4 Discussion

The primary contribution of this article is a method to derive and compute the transfer function for a size-structured population model in continuous time. This size-structured model is a leading example of a more general class of physiologically structured, continuous-time population models (PSPMs) in which individuals are classified by one or more continuously valued state variables (Metz and Diekmann, 1986; de Roos, 1997; Diekmann et al., 2007). We conjecture that the methods described here can be adapted to a wide variety of PSPMs.

Our study of the common Indo-Pacific coral *Pocillopora verrucosa* illustrates how transfer functions identify resonant oscillations that may emerge in suitably noisy environments. In the model, resonant oscillations are caused by intraspecific space competition in which high coral cover at the cycle peak interferes with the growth and recruitment of small, young colonies that will constitute the bulk of the population a few decades hence (Roughgarden et al., 1985; Pascual and Caswell, 1991; Artzy-Randrup et al., 2007). Resonant oscillations are more pronounced when colonies grow slowly (Fig. 4), when colony growth is only weakly affected by local crowding (Fig. S1), or when populations recruit primarily by larval immigration (Fig. S3). Because most coral populations today are severely degraded, it is unlikely that multi-decade resonance will be the chief dynamical feature of contemporary coral populations. Our model is thus offered to demonstrate how transfer functions can be derived and computed for PSPMs, and to suggest the potential for resonant oscillations in openly recruiting, size-structured populations with intraspecific space competition.

These methods point to a variety of avenues for future work. First, our coral example uses a simple description of how environmental variation impacts the vital rates that determine population dynamics. In particular, in the model, environmental fluctuations affect all individuals identically, regardless of their size. More mechanistic or more nuanced descriptions of the effects of environmental variation may lead to more complex transfer functions. Greenman & Benton have used network methods to show that population spectra in discrete-time models may depend strongly on which subset of the population is impacted by environmental fluctuations (Greenman and Benton, 2005*a*,*b*). Their results are based on analysis of the eigenstructure of the underlying model. It stands to reason that similar theory may hold for PSPMs, although general methods for calculating eigenvalues and eigenvectors for PSPMs are not yet known.

Second, like any linearization approach, transfer functions work best when environmental perturbations are modest and the dynamics remain in the neighborhood of a single attracting equilibrium. While the transfer function for the coral model accurately predicts the frequency and amplitude of stochastically excited oscillations under moderate environmental noise, we suspect the linearizaton works well in this case because the dynamics generated by the unforced model are relatively well behaved. In contrast, linearization fares less well at predicting spectral peaks in strongly nonlinear models capable of more complex dynamical behaviors (Blarer and Doebeli, 1999; Reuman et al., 2006). A more complete theory of frequency spectra in PSPMs that extends beyond the domain of locally linear approximations awaits future work.

Finally, the transfer-function approach used here envisions populations that are buffeted by environmental stochasticity. However, McKane and colleagues (McKane and Newman, 2005; Alonso et al., 2007; Black and McKane, 2010) have shown that resonant oscillations can be excited by demographic stochasticity as well. In their approach, the shape of the power spectra is predominantly (though not entirely) determined by the Jacobian of a fully stochastic model’s mean-field approximation. As a result, the power spectrum is largely governed by the system’s predominant modes of relaxation towards equilibrium, as those modes are captured by the eigenstructure of the linearized model (Greenman and Benton, 2005b). Thus, we anticipate that a fully stochastic version of a size-structured population model would exhibit similar resonance frequencies to those identified by the transfer function of a PSPM. A formal study of this possibility provides a promising direction for future research.

## 5 Appendix Computing the transfer function

To compute the transfer functions, we approximate the integrals in eq. 20 with a system of ODE that can be integrated forward in time (Kirkilionis et al., 2001). To begin, it is necessary to find the equilibrium values (*C*\*, *b*\*). While techniques for doing so appear in Kirkilionis et al. (2001), we include that calculation here for completeness. We seek to solve the system of equations

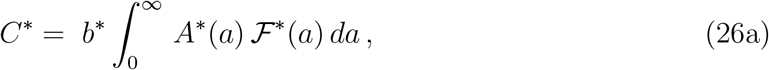

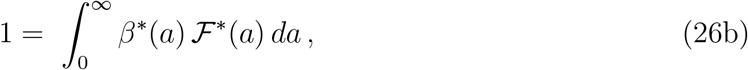

where we recall that *A*\*(*a*), 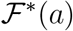, and *β*\*(*a*) all depend on *C*\*. Define the following integrals to age *a*:

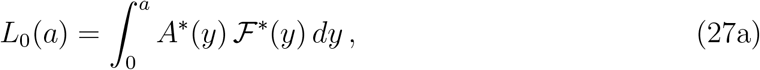

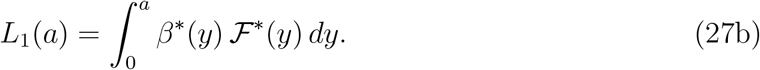

To avoid integrating to infinity, choose a large age, *a_ϵ_*, such that survival to age *a_ϵ_* is sufficiently small. For this article, we use *a_ϵ_* = 10^3^. We approximate *L_i_*(∞) by *L_i_*(*a_ϵ_*) for *i* = 0, 1. For a given guess of *C*\*, we can find *L_i_*(*a_ϵ_*) by integrating the following system of ODE

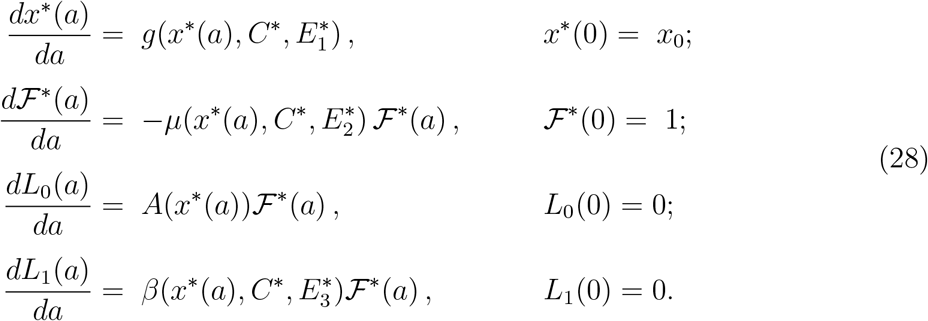

The pair (*C*\*, *b*\*) is then approximated by using a standard numerical root-finding algorithm to solve

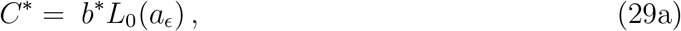

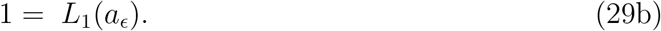

At each iteration of the root-finding algorithm, *L*_0_(*a_ϵ_*) and *L*_1_(*a_ϵ_*) are found by numerically solving eq. 28.

Having found (*C*\*, *b*\*), we then compute the transfer functions in a similar fashion. Define the following integrals to age *a*:

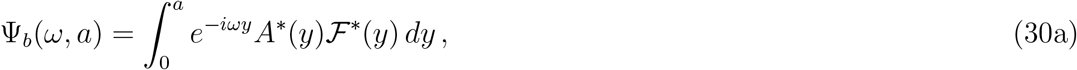

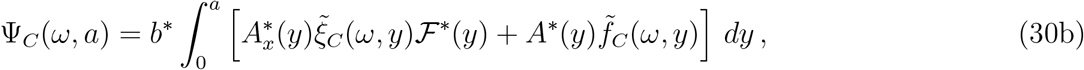

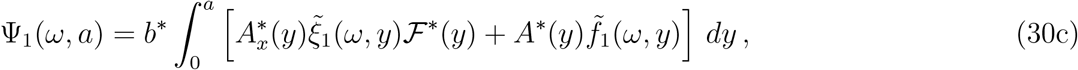

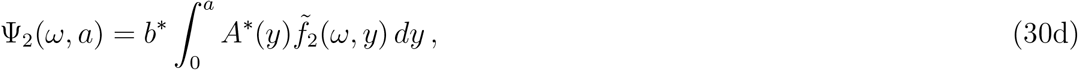

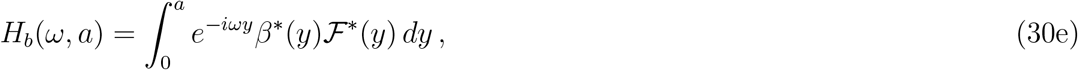

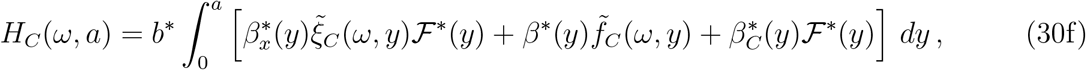

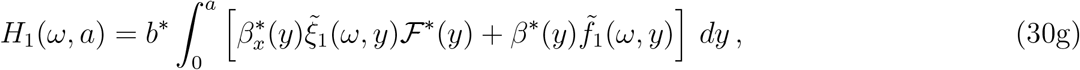

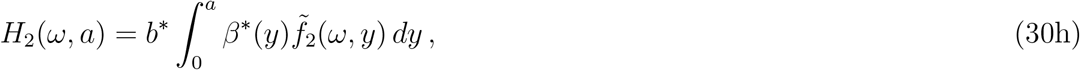

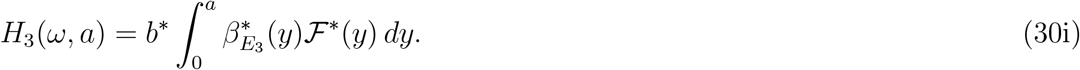

For a given *ω*, the integrals in eq. 30 can then be approximated by solving the following system of equations together with those in eq. 28 to age *a_ϵ_*:

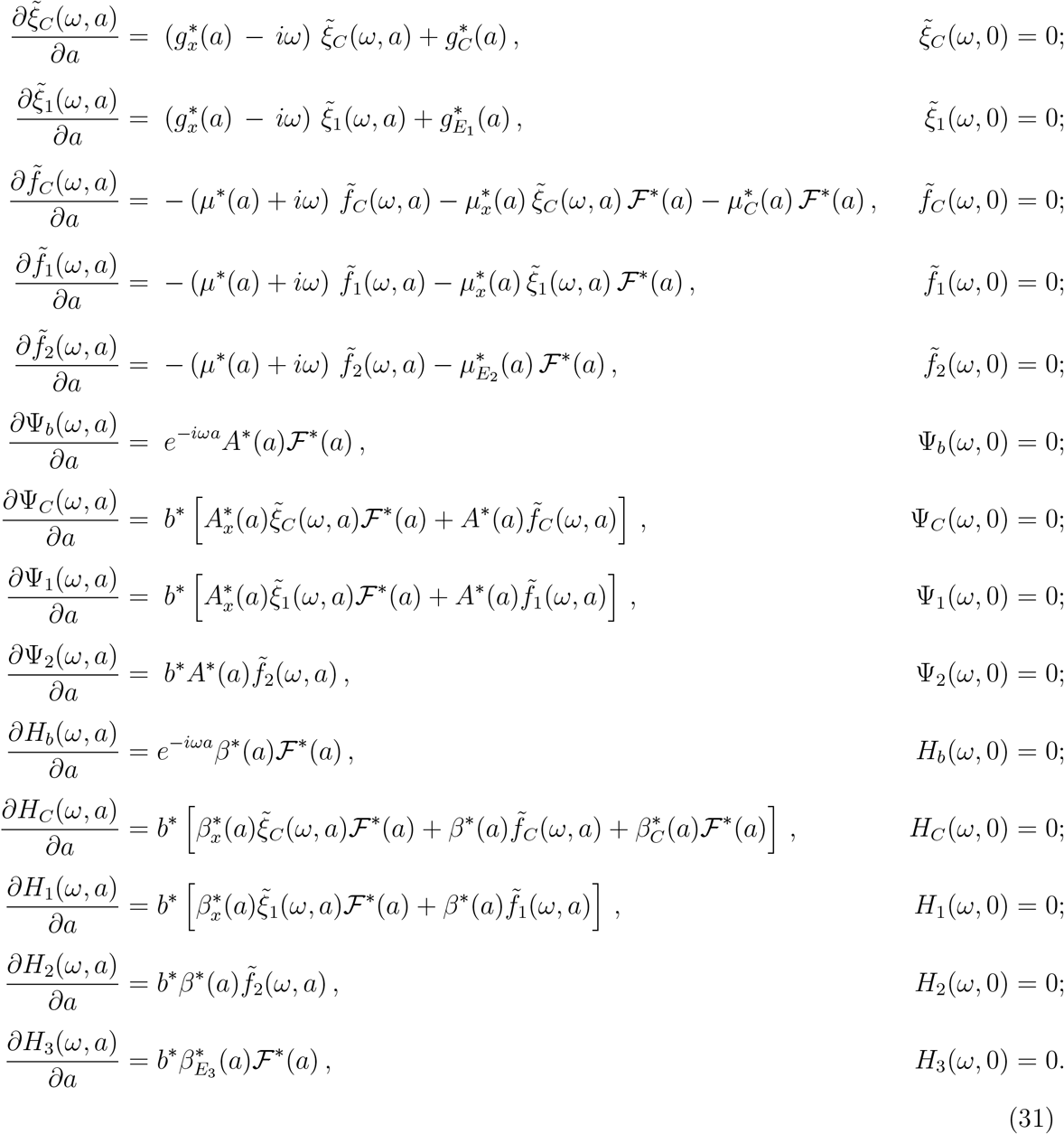

The transfer functions are then approximated by setting *ψ_b_*(*ω*) = Ψ_*b*_(*ω*, *a_ϵ_*); *ψ_C_* (*ω*) = Ψ*_C_* (*ω*, *a_ϵ_*); *ψ_i_*(*ω*) = Ψ_*i*_(*ω*, *a_ϵ_*) for *i* = 1, 2; *η_b_*(*ω*) = *H_b_*(*ω*, *a_ϵ_*); *η_C_* (*ω*) = *H_C_* (*ω*, *a_ϵ_*); and *η_j_*(*ω*) = *H_j_*(*ω*, *a_ε_*) for *j* = 1, 2, 3 in eq. 21.

## Acknowledgements

We thank Peter Edmumds, Bob Carpenter, and colleagues associated with the Mo’orea Coral Reef Long-Term Ecological Research project for their collaboration which inspired this work and for the *P. verrucosa* data used to parameterize our model. We also thank Marissa Baskett for her suggestion to explore resonance in these models, and we thank Matthew Spencer and anonymous reviewer for helpful comments that improved the article. This work was supported by NSF award OCE 14-15300 to KG, and by NSF support for the Mo’orea Coral Reef LTER.

## Supplementary Information

### S.1 Density-dependent growth

In this numerical experiment, we modified the component of the growth function that controls density dependence, *g*_1_(*C*(*t*)). First, we tuned the average larval immigration rate to yield an equilibrium coral density under constant environments of exactly *C*\* = 1/2. We then defined the density-dependent component of growth as

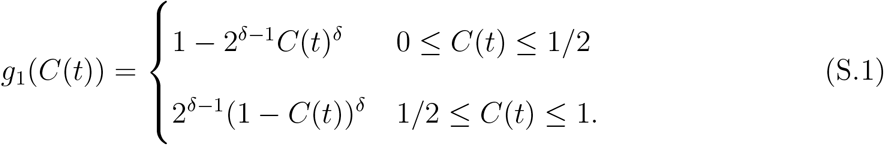

In eq. S.1, the strength of density dependence is determined by the parameter *δ* (Fig. S1a). A value of *δ* = 1 corresponds to growth being proportional to free space, *g*_1_(*C*(*t*)) = 1 − *C*(*t*). A value of *δ* > 1 corresponds to strong density dependence, in which growth drops rapidly when cover is in the neighborhood of *C*\*. Conversely, *δ* < 1 corresponds to weak density dependence, in which growth is less affected by coral cover in the neighborhood of *C*\*. We then computed the resonant frequency and modulus of the transfer function at the resonant frequency for various values of *δ*. These computations show that resonant cycles become more frequent and smaller in amplitude as density dependence becomes stronger (Fig. S1B,C). Sample dynamics are shown in Fig. S2.

### S.2 Internal vs. external recruitment

This numerical experiment investigates how the blend of internal vs. external recruitment affects resonant oscillations. In this experiment, *p* ∈ [0, 1] gives the proportion of recruits that arise from internal reproduction. For internal recruitment, we assume that each colony produces larvae in proportion to its planar area, and that internal reproduction is multiplied by the same environmental forcing that impacts external recruitment, *E*_3_(*t*). Thus, colonies in the population contribute recruits at a rate *pγE*_3_(*t*)*A*(*x*)(1 − *C*(*t*)), where *γ* is tuned so that the equilibrium coral cover *C*^⋆^ is independent of *p*. The total effective reproduction for each colony is then

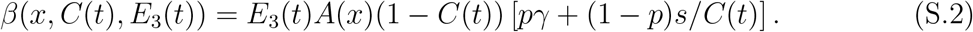

**Figure S1:**
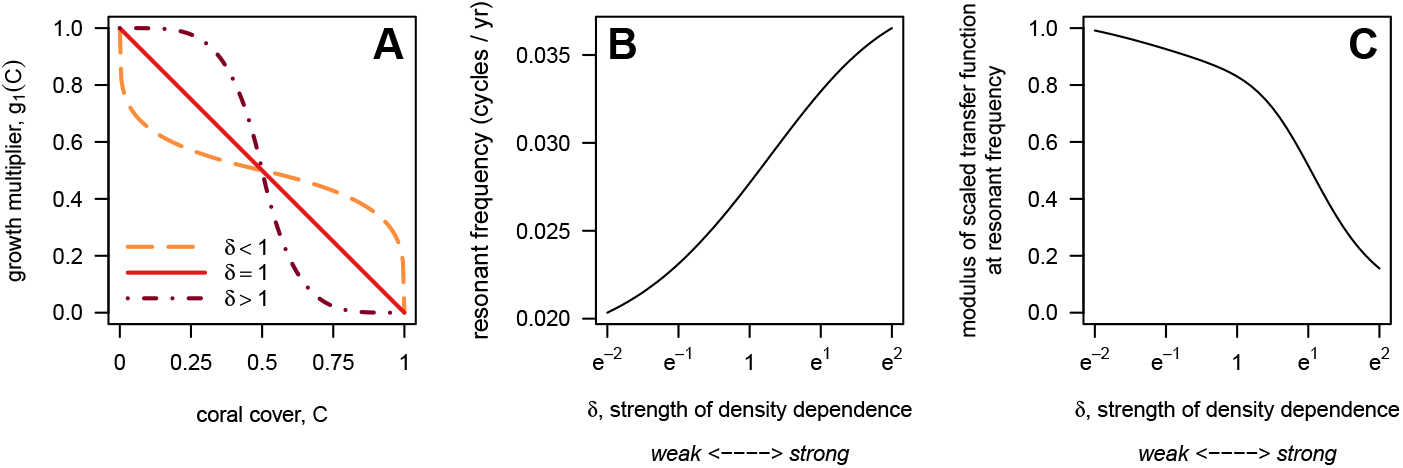
Increasing the strength of density dependence in colony growth increases the frequency and decreases the strength of population resonance. (A): The density-dependent component of growth, *g*_1_(*C*(*t*)), plotted vs. coral cover *C*(*t*) for three example values of *δ*. (B): The resonant frequency for the scaled transfer function for recruitment (*T*_3,*s*_(*ω*)) and (C) the modulus of this transfer function at the resonant frequency vs. the strength of density dependence, as captured by *δ*. Similar relationships hold for the transfer functions for growth (*T*_1*,s*_(*ω*)) and mortality (*T*_2,*s*_(*ω*)).

**Figure S2:**
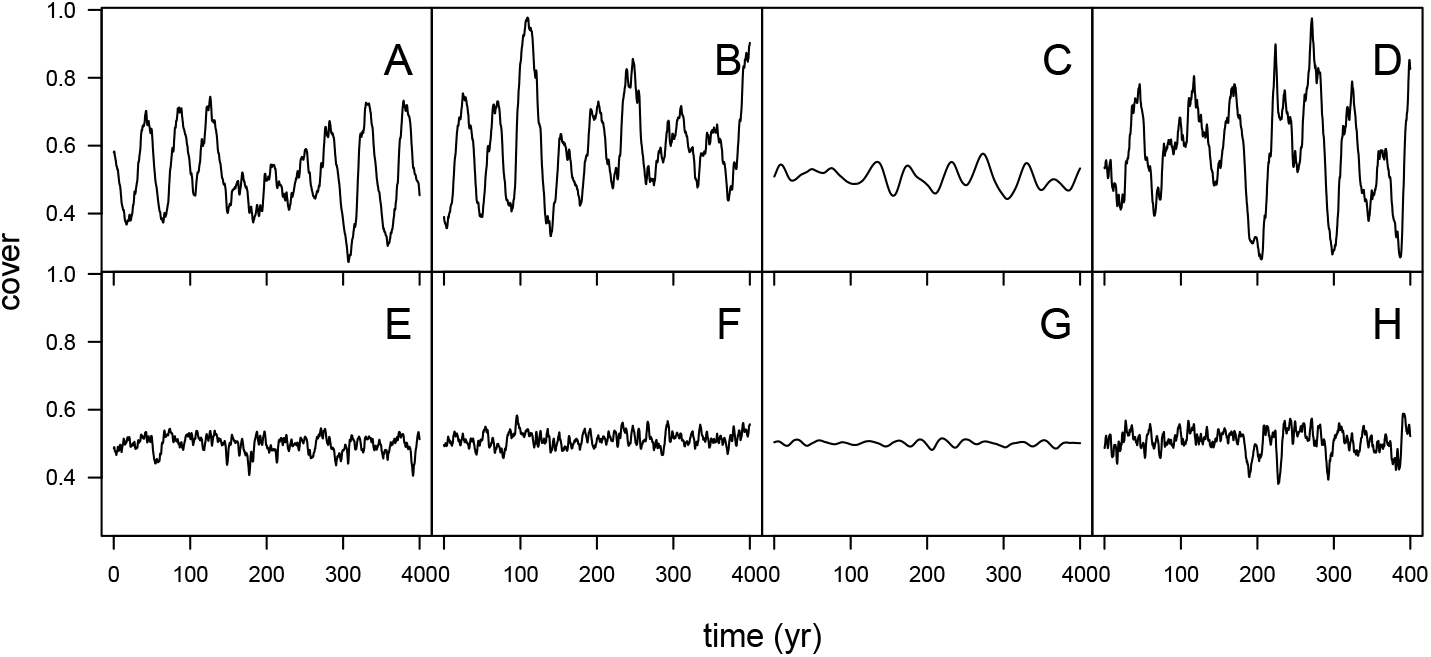
The strength of density dependence affects population resonance in the *P. verrucosa* model. Each panel shows 400 yr of simulated dynamics. The environmental forcing variable(s) is (are) either (A, E) growth, (B, F) mortality, (C, G) recruitment, or (D, H) growth, mortality, and recruitment together. The top row (A – D) shows weak density dependence in growth (*δ* = *e*^−1.5^, shown in the dashed curve in Fig. S1A), and the bottom row (E – H) shows strong density-dependence in growth (*δ* = *e*^1.5^, shown in the dot-dashed curve in Fig. S1A). Simulations were obtained using the same methods as Fig. 1.

**Figure S3:**
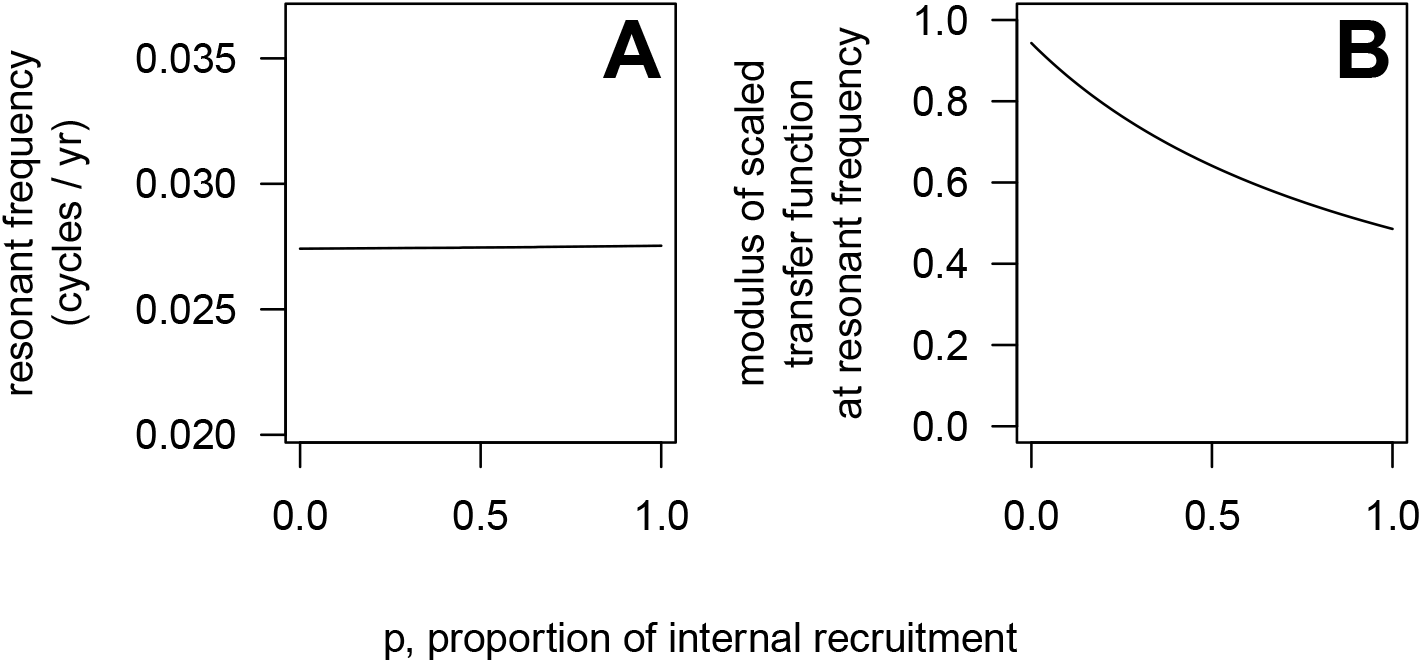
The relative blend of internal vs. external recruitment has little effect on population resonance in the *P. verrucosa* model. (A) Resonant frequency and (B) modulus at the resonant frequency of the scaled transfer function for recruitment (*T*_3*,s*_(*ω*)) vs. *p*, the proportion of internal recruitment.

We then varied *p* and computed the resonant frequency and modulus of the transfer function at the resonant frequency using the scaled version of *T*_3_(*ω*). The resonant frequency of the population is almost completely unaffected by whether the population is open or closed (Fig. S3A). In contrast, the amplitude of resonant oscillations is greatest when recruits come from outside the population and lowest when recruits are produced locally (Fig. S3B).

